# The conserved QTQTX motif in the SARS-CoV-2 spike protein is dispensable for cleavage and lung cell entry of the emerging variant BA.3.2

**DOI:** 10.64898/2026.05.04.722688

**Authors:** Lu Zhang, Nianzhen Chen, Amy Eichmann, Inga Nehlmeier, Ryan Hisner, Christine Happle, Georg M. N. Behrens, Markus Hoffmann, Stefan Pöhlmann

**Affiliations:** Infection Biology Unit, German Primate Center – Leibniz Institute for Primate Research, Göttingen, Germany; Faculty of Biology and Psychology, Georg-August University Göttingen, Göttingen, Germany; Institute of Molecular Virology, Ulm University Medical Center, 89081 Ulm, Germany; Division of Computational Biology, Department of Integrative Biomedical Sciences, Institute of Infectious Diseases and Molecular Medicine, University of Cape Town, Cape Town, South Africa; Department for Rheumatology and Immunology, Hannover Medical School, Hannover, Germany; German Centre for Lung Research (DZL), partner site Hannover BREATH, Hannover, Germany; German Centre for Infection Research (DZIF), partner site Hannover-Braunschweig, Hannover, Germany; Excellence Cluster RESIST (EXC 2155), Hannover Medical School, Hannover, Germany; German Center for Infection Research (DZIF), associated partner site Göttingen, Göttingen, Germany

## Abstract

The furin motif in the SARS-CoV-2 spike (S) protein is important for lung cell entry. It is embedded in an extended loop structure and preceded by a highly conserved QTQTX motif that is required for efficient furin cleavage of the SARS-CoV-2 WA-1 S protein. BA.3.2 is an emerging SARS-CoV-2 saltation variant that is spreading globally in April 2026 and encodes for a highly mutated S protein. Here, we analyzed whether the QTQTX motif is also required for spike protein cleavage and lung cell entry of BA.3.2. We report that two patient-derived spike sequences of the BA.3.2 subvariant BA.3.2.2 lack the first QT repeat of the QTQTX motif and show that this motif is largely dispensable for both cleavage and lung cell entry of BA.3.2.2, which we found to depend on TMPRSS2. Our results suggest that the reconfiguration of the BA.3.2 S protein during persistent infection may have significantly altered the determinants of furin cleavage.

**IMPORTANCE:** The furin motif in the SARS-CoV-2 spike (S) protein is unique among sarbecoviruses and constitutes a virulence determinant. A QTQTX motif located immediately upstream of the furin motif is required for furin cleavage of the S protein of the virus that circulated early in the pandemic. Here, we show that the QTQTX motif is largely dispensable for S protein processing and S protein-driven lung cell entry of the emerging saltation variant BA.3.2, which is currently spreading globally. Thus, BA.3.2 evolution within immunocompromised individuals may have relaxed the requirements for furin processing of the spike protein.

## INTRODUCTION

The spike (S) protein of SARS-CoV-2 facilitates viral entry into host cells and is the key target of the neutralizing antibody response (1, 2). For entry, the surface unit, S1, of the S protein binds to the cellular receptor, angiotensin-converting enzyme 2 (ACE2) (1, 2), and the transmembrane unit, S2, fuses the viral with a cellular membrane, allowing delivery of the viral genome into the cellular cytoplasm. Membrane fusion requires cleavage of the S protein by host cell proteases (3). For entry into lung cells, the S protein is first cleaved at the S1/S2 site located in a protruding loop at the border between the S1 and S2 subunit. The cleavage site is defined by a furin motif that is cleaved by the proprotein convertase furin or related proteases in the constitutive secretory pathway of infected cells (4–6). Subsequently, the S protein is processed at the S2’ site by the cell surface protease TMPRSS2 during entry into lung cells (4, 5). The S2’ site is located within the S2 subunit and its cleavage is believed to trigger or at least accelerate the membrane fusion reaction (7).

The presence of a furin motif in the SARS-CoV-2 S protein is unique among sarbecoviruses and is the central determinant of furin cleavage and a virulence factor (8, 9). Further, it is required for efficient transmission (9, 10). However, a previous study showed that a highly conserved QTQTX motif directly preceding the furin motif is also important for furin cleavage, lung cell entry and virulence in the context of SARS-CoV-2 variant circulating in the early phase of the pandemic (11). It is presumed that the QTQTX motif is required for exposure of the furin motif within the cleavage loop, making it readily accessible to furin (11). However, these findings were made with SARS-CoV-2 WA-1 and it was not examined whether the S proteins of other SARS-CoV-2 variants also depend on the QTQTX motif for furin cleavage and lung cell entry.

BA.3.2 is a saltation variant and it is believed to have evolved in a persistently infected, immunocompromised individual and subsequently entered the SARS-CoV-2 circulation (12). It was first detected in November 2024 in South Africa and subsequently spread to Australia, Europe, North America and several Asian countries (12, 13). BA.3.2 dominates in several European countries and parts of Australia as of March 2026 but dominance is not universal and spread of BA.3.2 is generally slower than that observed for previous variants. As compared to the parental BA.3 lineage, the BA.3.2 S protein contains 53 mutations, including mutations and deletions in the N-terminal domain antigenic supersite and epitope altering mutations in the receptor binding domain (14). These mutations are compatible with robust entry into several cell lines but ACE2 binding and lung cell entry are diminished (13, 15).

Since its emergence, BA.3.2 has further evolved and diverged into two main subvariants, BA.3.2.1 and BA.3.2.2, which themselves evolved and diverged into their own subvariants, such as RD.* (diverged from BA.3.2.1) and RE.* (diverged from BA.3.2.2) (**Supplementary Fig. S1**). We found that two RE.1.1 sequences from Western Australia, collected on October 4, 2025 (EPI_ISL_20221586), and November 4, 2025 (EPI_ISL_20244228), harbored S proteins with deletion in the QTQTX motif, which removed the first QT repeat (**Fig. 1A**). Since the S proteins of BA.3.2.2 and RE.1.1 are identical on the amino acid level, we examined the role of the QTQTX motif in BA.3.2.2 S protein cleavage and lung cell entry.

**FIG 1.**
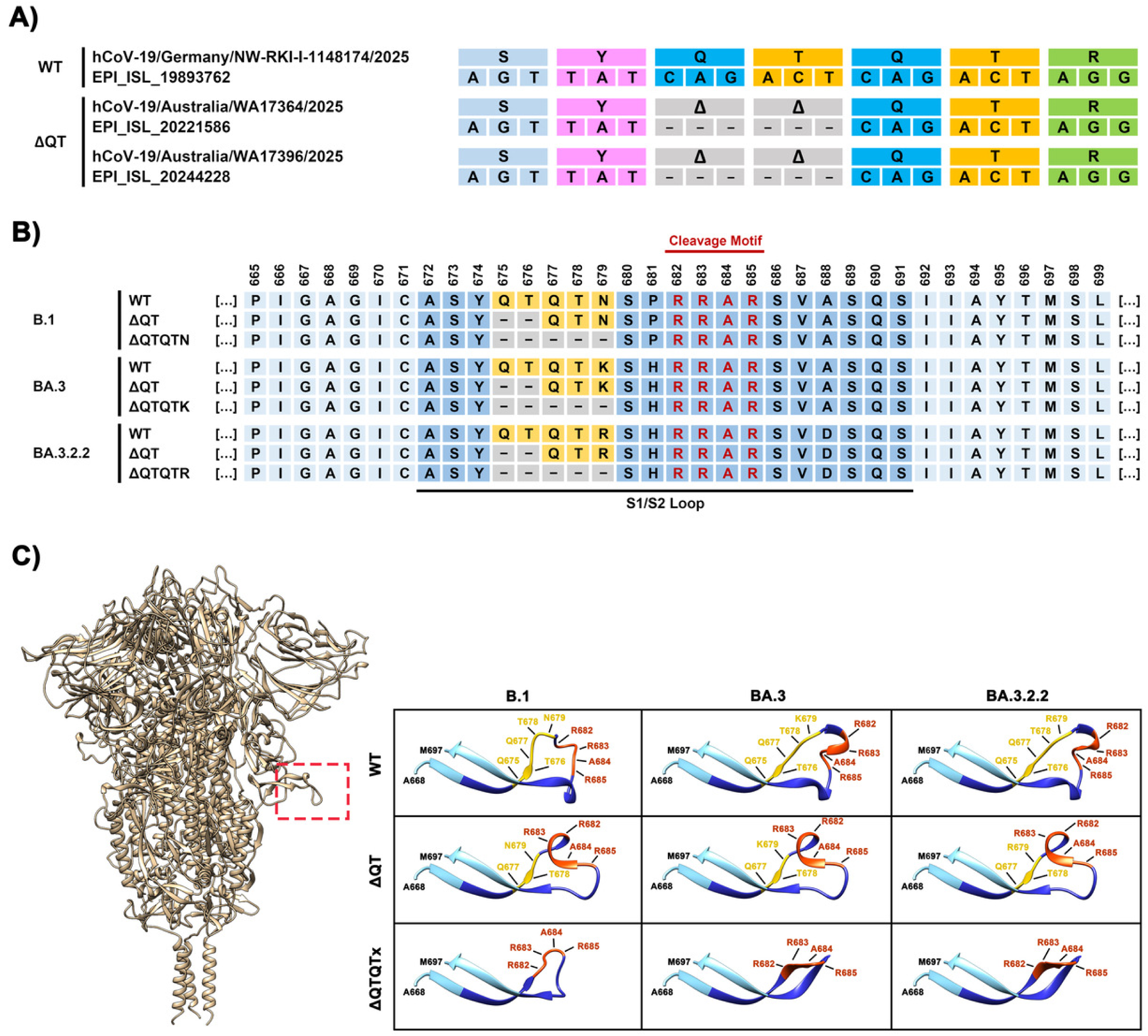
Naturally occurring and engineered deletions in the QTQT motif. **(A)** Deletions in the QTQTx motif observed in two individuals infected with the BA.3.2.2 variant in Australia. **(B)** Engineered deletions in the QTQTx motif analyzed within the present study. **(C)** Protein structure of the SARS-CoV-2 S protein (based on PDB:6XR8 ((35)) with the S1/S2 loop highlighted (left) and S1/S2 loop structures predicted by homology modelling (right). The amino acid numbering is based on B.1 S protein wild type.

We report that the QTQTX motif is required for cleavage of the SARS-CoV-2 B.1 (i.e. Wuhan-01 + D614G) and BA.3 S proteins but is largely dispensable for cleavage of the BA.3.2.2 S protein. In agreement with this finding, an intact QTQTX motif was essential for robust lung cell entry driven by the B.1 and BA.3 but not the BA.3.2.2 S protein.

## RESULTS

### The QTQTX motif is dispensable for BA.3.2.2 S protein cleavage

In order to analyze the role of the QTQTX motif in proteolytic processing of the BA.3.2.2 S protein and S protein-driven cell entry, we deleted either the first QT repeat or the entire QTQTX motif in the S proteins of SARS-CoV-2 B.1 (B.1 = Wuhan-1 plus D614G), BA.3 and the BA.3.2 subvariant BA.3.2.2 (**Fig. 1B**).

The S proteins of B.1, BA.3 and BA.3.2.2 differ in their respective QTQTX motif by the last amino acid residue, which is either an asparagine (N, B.1), a lysine (K, BA.3) or an arginine (R, BA.3.2.2) (**Fig. 1B**). Protein structure prediction by homology modelling with Swiss Model (https://swissmodel.expasy.org/) (16) indicated minor differences between the S1/S2 loop structures of wildtype (WT) B.1, BA.3 and BA.3.2.2 S proteins that are mainly linked to the N-terminal part of the S1/S2 cleavage site. Deletion of the first QT repeat was compatible with continued exposure of the S1/S2 site while deletion of the entire QTQTX motif resulted in major structural rearrangements within the respective S1/S2 loop structures that were associated with profound changes in the exposure of the S1/S2 cleavage site (**Fig. 1C**). Immunoblot analysis of transfected 293T cells revealed that deletion of the first QT repeat reduced cleavage of B.1 S protein but had no effect on cleavage of the BA.3 and the BA.3.2.2 S proteins (**Fig. 2A and 2B**). Deletion of the entire QTQTX motif markedly reduced cleavage of the B.1 and BA.3 S proteins but barely diminished cleavage of the BA.3.2.2 S protein (**Fig. 2A and 2B**). Thus, the QTQTX motif is required for cleavage of WT S protein, in agreement with a previous study (11), and the BA.3 S protein, but is largely dispensable for cleavage of the highly mutated BA.3.2.2 S protein.

**FIG 2.**
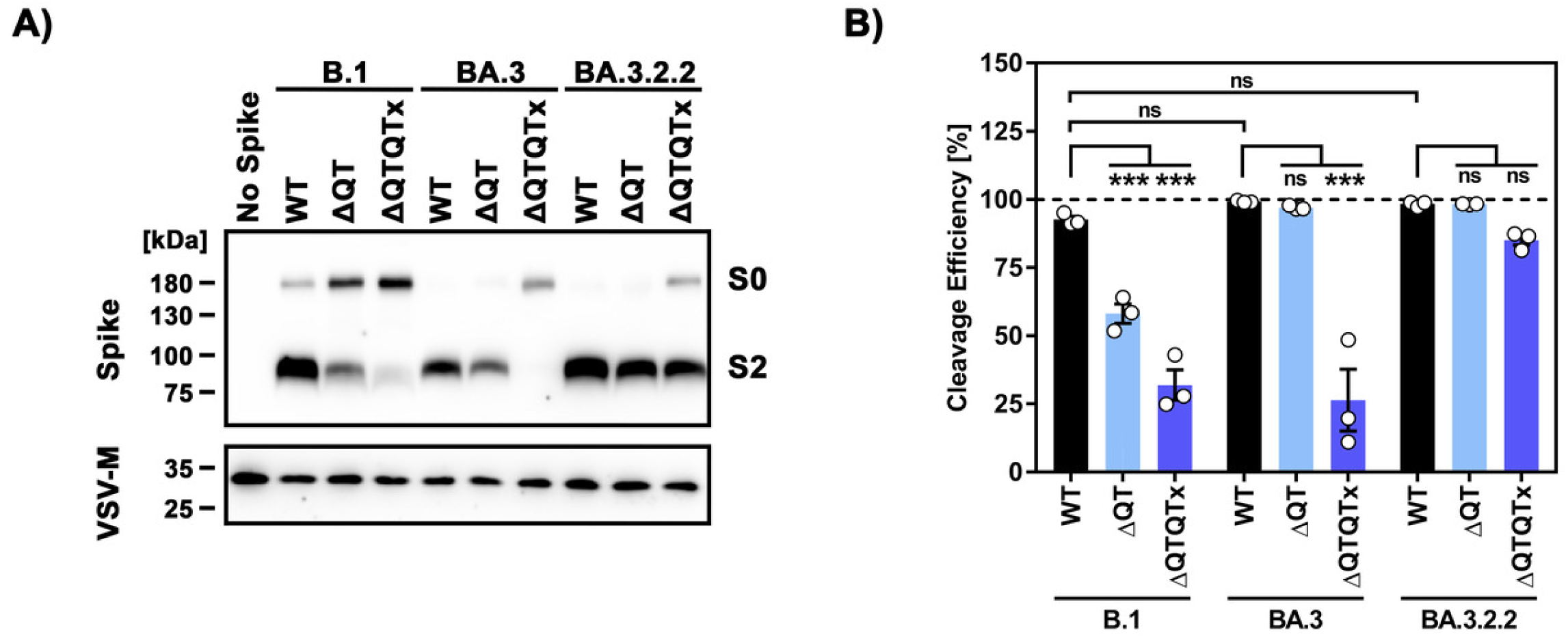
The QTQTX motif is dispensable for cleavage of the BA.3.2.2 S protein. (A) Equal volumes of pseudoviruses bearing the indicated S proteins were subjected to SDS-PAGE followed by immunoblot analysis to assess S protein incorporation into viral particles and processing at the S1/S2 cleavage site. Immunodetection was performed using an antibody specific for the SARS-CoV-2 S protein and a peroxidase-conjugated secondary antibody. Detection of vesicular stomatitis virus matrix protein (VSV-M) served as a loading control. A representative immunoblot is shown, the results were confirmed in two additional independent biological replicates. (B) Band intensities of the S0 and S2 species were quantified by densitometry using ImageJ. For each sample, the combined S protein signal (S0 + S2) was set to 1, and the relative abundance of S0 and S2 is shown as the mean of three independent biological replicates. Statistical significance was asessed by one-way analysis of variance (ANOVA) with Tukey’s multiple comparison test (not significant [ns], p >0.05; *, p ≤0.05; **, p ≤0.01; ***, p ≤0.001).

### The deletion of the QTQTX motif does not diminish BA.3.2.2 S protein-driven cell-cell fusion

We next investigated the role of the QTQTX motif in S protein-driven cell-cell fusion, employing a previously reported cell-cell fusion assay based on 293T effector cells expressing S protein and 293T control target cells overexpressing ACE2 alone or in combination with TMPRSS2 (17). We found that cell-cell fusion driven by all S proteins was more robust when TMPRSS2 was coexpressed with ACE2 on target cells, while fusion driven by Nipah virus fusion protein and glycoprotein (NiV-F+G) remained unaffected by the presence of TMPRSS2, as expected (5) (**Fig. 3A**). Deletion of the first QT repeat markedly reduced fusion with both target cells driven by the B.1 S protein, while deletion of the first QT repeat in the S proteins of BA.3 and BA.3.2.2 S proteins had only a minor or no impact on cell-cell fusion (**Fig. 3A**), in keeping with the S protein cleavage data (**Fig. 2**). Deletion of the entire QTQTX motif further reduced B.1 S protein-driven fusion with target cells expressing ACE2 but only slightly reduced fusion with cells coexpressing ACE2 and TMPRSS2 (**Fig. 3A**). Similarly, deletion of QTQTX motif markedly reduced BA.3 S protein-mediated fusion with ACE2 expressing cells but had only a moderate negative effect on fusion with ACE2 and TMPRSS2 coexpressing target cells. Finally, the absence of the QTQTX motif did not reduce fusion with ACE2 expressing or ACE2 and TMPRSS2 coexpressing cells driven by the BA.3.2.2 S protein (**Fig. 3A**). Thus, the QTQTX motif is dispensable for BA.3.2.2 S protein-mediated cell-cell fusion but is required for fusion mediated by B.1 and BA.3 spike proteins.

**FIG 3.**
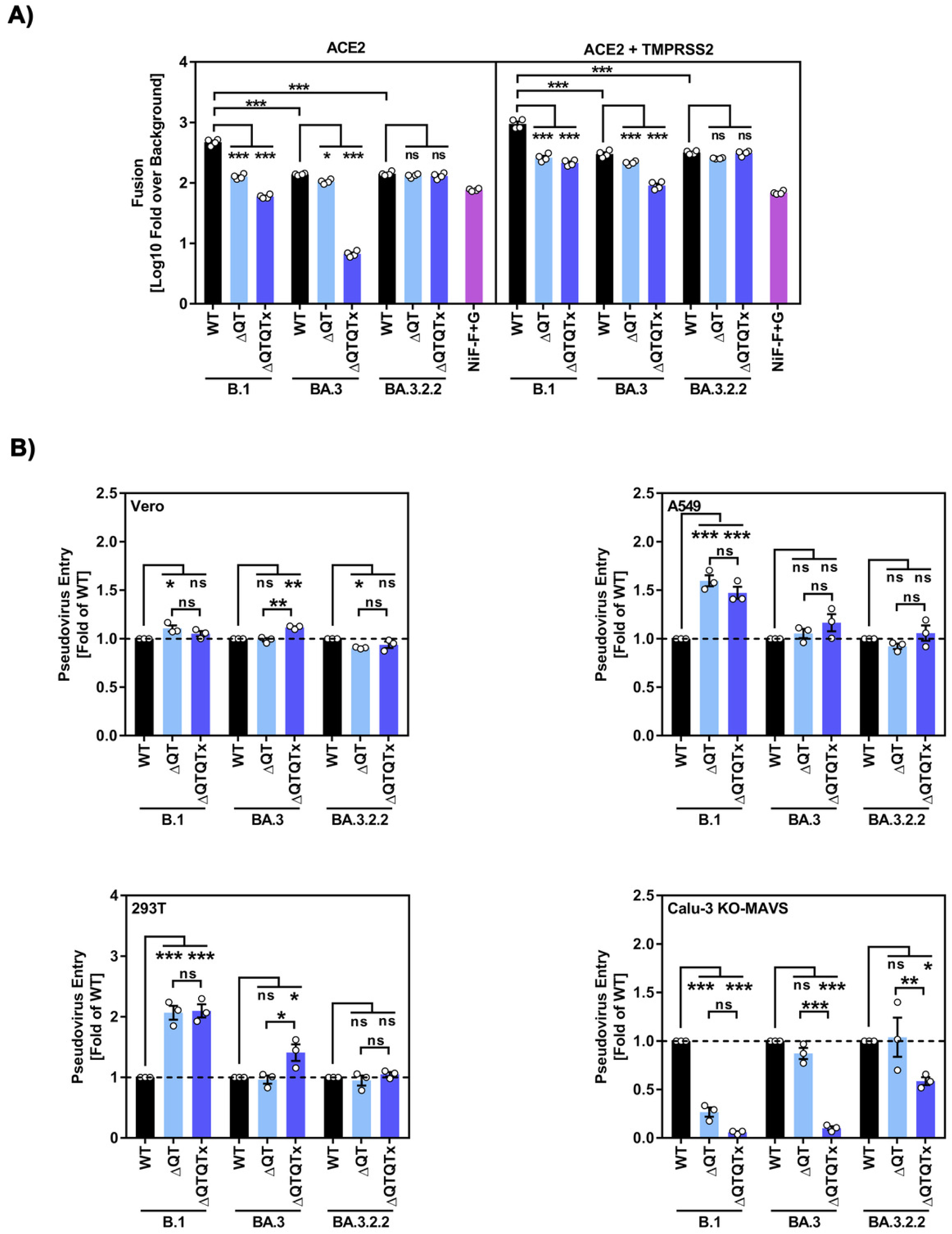
Deletion of the QTQTX motif does not reduce cell-cell and virus-cell fusion driven by the BA.3.2.2 S protein. 293T effector cells co-expressing the beta-galactosidase alpha fragment and the indicated S proteins or Nipah virus fusion- and glycoprotein (NiV-F+G), or no viral glycoprotein (control effector cells) were co-cultured with 293T target cells coexpressing the beta-galactosidase omega fragment and either ACE2 or ACE2 and TMPRSS2. Efficiency of cell-cell fusion was determined by quantification of beta-galactosidase activity in cell lysates and data were normalized against the signals obtained upon co-culture of target cells with control effector cells expressing no viral glycoprotein. The mean of four biological replicates (each conducted with four technical replicates) is shown. Error bars represent the SEM. (B) Pseudoviruses bearing the indicated S proteins were used to infect different target cell lines. Entry efficiency was determined 16 hours post infection by quantification of pseudovirus-expressed firefly luciferase in cell lysates. The results were normalized against signals obtained for pseudoviruses equipped with B.1 S protein (set as 1). Data show the mean of three biological replicates performed with four technical replicates, error bars indiate SEM. Statistical significance was asessed by one-way ANOVA with Tukey’s multiple comparison test (ns, p >0.05; *, p ≤0.05; **, p ≤0.01; ***, p ≤0.001).

### The QTQTX motif is largely dispensable for BA.3.2.2 S protein-mediated virus-cell fusion

In order to examine the role of the QTQTX motif in virus-cell fusion, we employed vesicular stomatitis virus (VSV)-based pseudotypes, which adequately model key aspects of SARS-CoV-2 entry into cells and its inhibition (18). As target cells, we chose Calu-3 lung cells, which allow for TMPRSS2-dependent entry (1), which in turn requires furin cleavage (5), and A549 lung and 293T and Vero kidney cells, which allow for furin-independent, cathepsin L-dependent entry (1).

Specifically, a Calu-3 cell subline lacking MAVS (KO-MAVS, (19)) was chosen since the inactivation of MAVS does not alter the entry mechanism but allows for more robust infection with VSV-based pseudotypes, most likely due to less efficient inhibition of VSV by components of the interferon system.

We found that deletion of the first QT repeat and, particularly, the entire QTQTX motif markedly reduced Calu-3 KO-MAVS lung cell entry of pseudoparticles (pp) bearing the B.1 S protein (B.1_pp_) (**Fig. 3B and Supplemental Fig. S2**). In contrast, Calu-3 MAVS-KO lung cell entry of pseudotypes bearing the BA.3 or BA.3.2.2 S proteins (BA.3_pp_, BA.3.2_pp_) was not affected by deletion of the first QT repeat, and deletion of the entire QTQTX motif markedly reduced BA.3_pp_ but only modestly reduced BA.3.2.2_pp_ cell entry (**Fig. 3B**). Finally, deletion of the first QT repeat or the entire QTQTX motif had no effect or increased B.1_pp_, BA3_pp_ and BA.3.2.2_pp_ entry into A549, Vero and 293T cells. The latter finding is in agreement with the notion that propagation of SARS-CoV-2 in cells that allow only for cathepsin L-dependent entry can result in rapid loss of the furin cleavage site and adjacent sequences, which increases cell entry (20, 21). In sum, the QTQTX motif is required for robust lung cell entry of B.1_pp_ and BA.3_pp_ but largely dispensable for BA3.2.2_pp_.

### BA.3.2_pp_ entry into Calu-3 lung cells is inhibited by Camostat

We next probed protease dependence of BA.3_pp_ and BA.3.2.2_pp_ entry. For this, we employed Camostat, a serine protease inhibitor active against TMPRSS2, and MDL28170, an inhibitor of cysteine proteases including cathepsin L (1). Vero cells were used as targets since they are well documented to allow exclusively for cathepsin L-dependent entry (1). Inhibition of Vero cell entry by protease inhibitors was compared to inhibition of Calu-3 KO-MAVS lung cell entry.

We found that entry of all pseudotypes into Vero cells was efficiently blocked by MDL28170 (**Fig. 4A, upper panel**), in agreement with expectations (1). It was noted that inhibition of B.1_pp_ entry was more efficient than that of BA.3_pp_ and BA.3.2.2_pp_ entry, as expected in light of the higher efficiency of cathepsin L usage by Omicron subvariants as compared to B.1_pp_. Further, Camostat efficiently inhibited entry of B.1_pp_, BA.3_pp_ and BA.3.2.2_pp_ into Calu-3 KO-MAVS cells (**Fig. 4A, lower panel**), with inhibition of BA.3_pp_ being somewhat less robust than that of B.1_pp_ and particularly BA.3.2.2_pp_. These results suggest that BA.3 and BA.3.2.2 join BA.2.86 (17, 22) as the only Omicron subvariants that employ TMPRSS2 for Calu-3 lung cell entry.

**FIG 4.**
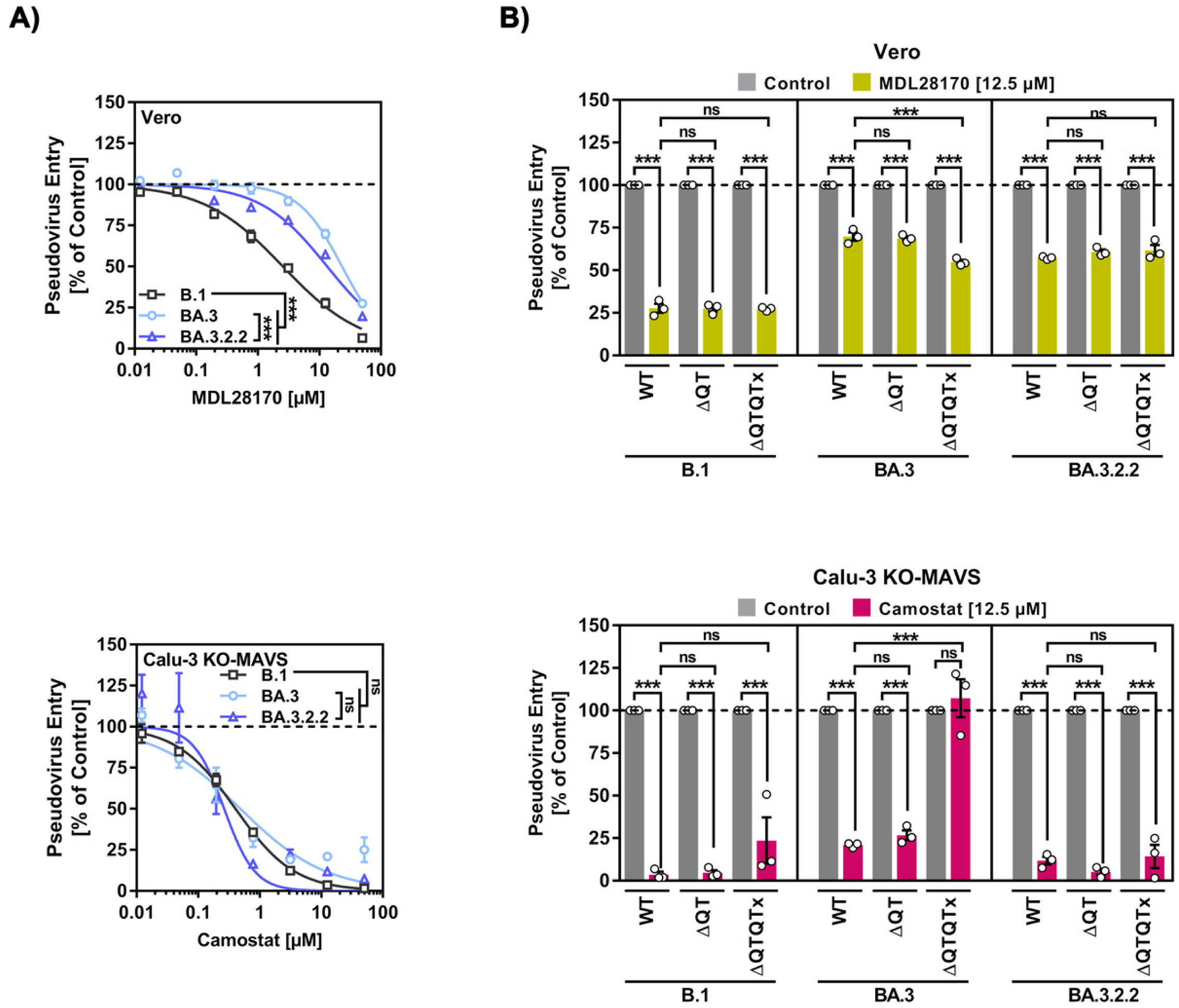
Calu-3 cell entry is Camostat-sensitive. (A) Vero and Calu-3 target cells were preincubated for 30 minutes with the indicated inhibitors followed by inoculation with pseudoviruses bearing the indicated S proteins and further incubation in the presence of inhibitor for 16 hours. Entry efficiency was determined by quantification of pseudovirus-expressed firefly luciferase in cell lysates and normalized against signals obtained in the absence of inhibitor (set as 100%). Data show the mean of three biological replicates performed with four technical replicates, error bars indiate SEM. Statistical significance was asessed by two-way ANOVA with Tukey’s multiple comparison test (ns, p >0.05; *, p ≤0.05; **, p ≤0.01; ***, p ≤0.001). **(B)** The experiment was conducted as describe for panel A with the exeptions that fixed concentrations of inhibitor were used (12.5 µM MDL28170; 50 µM Camostat) and particles bearing wildtype (WT) or mutant S proteins were investigated. Data show the mean of three biological replicates performed with four technical replicates, error bars indiate SEM. Statistical significance was asessed by one-way ANOVA with Tukey’s multiple comparison test (ns, p >0.05; *, p ≤0.05; **, p ≤0.01; ***, p ≤0.001).

Finally, we investigated whether deletion of the first QT repeat or the entire QTQTX motif impacts protease dependence, using fixed concentrations of MDL28170 (12.5 µM) and Camostat (50 µM). Our results obtained with Vero cells confirmed the reduced MDL28170-sensitivity of BA.3_pp_ and BA.3.2.2_pp_ as compared to B.1_pp_, that we had noted above, and showed that deletion of the first QT repeat or the entire QTQTX motif did not appreciably alter MDL28170 sensitivity (**Fig. 4B, upper panel**). Inhibition of Calu-3 KO-MAVS entry by Camostat confirmed that Camostat-sensitivity of BA.3_pp_ was slightly lower than that B.1_pp_ and BA.3.2.2_pp_ and showed that deletion of the first QT repeat or the entire QTQTX motif had little impact on Camostat-sensitivity of BA.3.2.2_pp_, although a slight Camostat resistance was observed for mutant ΔQTQTX (**Fig. 4B, lower panel)**. In contrast, deletion of the QTQTX motif rendered Calu-3 KO-MAVS cell entry of B.1_pp_ partially and that of BA.3_pp_ fully Camostat-insensitive, suggesting that the residual Calu-3 KO-MAVS cell entry of these mutants (**Fig. 3B**) is at least partially TMPRSS2-independent, in keeping with the concept that absence of pre-cleavage at the S1/S2 sites skews entry from TMPRSS2 towards cathepsin L dependence (5).

## DISCUSSION

The integrity of the furin motif is essential for SARS-CoV-2 S protein processing at the S1/S2 site in infected cells and for viral entry into cultured lung cells (1, 4). In agreement with these findings, SARS-CoV-2 without the furin motif exhibits reduced replication in the respiratory tract in rodent models and infection is associated with reduced pathology (8, 9). Further, mutation of the furin motif reduces but may not abrogate transmission (9, 10). The QTQTX motif in the SARS-CoV-2 S protein, which is absent from most sarbecoviruses excluding some bat and pangolin-derived RBD clade 1b sarbecoviruses (e.g. RaTG13) (**Supplementary Fig. S3**), is also important for efficient processing of the S protein, infection of Calu-3 human lung cells and full viral pathogenesis in a Syrian hamster model (11). However, these findings were made with SARS-CoV-2 circulating early in the pandemic (SARS-CoV-2 WA-1) and it is unknown whether later variants also depend on the QTQTX motif. Here, we provide evidence that the QTQTX motif is largely dispensable for S protein cleavage and lung cell entry of the emerging BA.3.2 variant.

The QTQTX motif present in SARS-CoV-2 circulating early in the pandemic is believed to be required for the flexibility of the cleavage loop and thus the accessibility of the furin cleavage motif to furin and other proprotein convertases, which ensures efficient S protein cleavage (11). Counterintuitively, the deletion of the QTQTX motif was compatible with full viral replication in the hamster respiratory tract but not with full viral pathogenesis, for at present unclear reasons (11). One can speculate that either species specific mutations in furin or potentially higher expression of furin in the hamster as compared to the human respiratory tract might have allowed for efficient replication of the SARS-CoV-2 delta QTQTX variant in hamsters. If so, one would need to also postulate that the reduced pathogenesis despite unabated viral replication resulted from a previously unappreciated role of the QTQTX motif in viral pathogenesis. A role of the QTQTX motif in SARS-CoV-2 infection beyond furin cleavage is supported by the finding that mutating the threonine residues within QTQTX abrogates O-glycosylation and reduces TMPRSS2-dependent host cell entry but does not impact S protein processing by furin (11). However, the underlying mechanism is unknown. Collectively, the QTQTX motif can promote S protein cleavage at the S1/S2 site by furin, by increasing the structural flexibility of the cleavage loop, and promotes TMPRSS2-dependent entry and COVID-19 pathogenesis, via so far unknown mechanisms.

The markedly reduced Calu-3 cell entry driven by the B.1 S protein lacking either the first QT repeat or the entire QTQTX motif confirmed the data published for SARS-CoV-2 WA-1 (11). This was expected, since both S proteins are highly similar. A comparable tendency was observed for BA.3_pp_, although Calu-3 cell entry driven by this S protein was less dependent on the first QT repeat as compared to the B.1 S protein, potentially reflecting subtle differences in S protein cleavage efficiency. The robust Calu-3 lung cell entry of BA.3.2.2_pp_ observed upon deletion of the QTQTX motif was in agreement with the efficient cleavage of this S protein mutant. Calu-3 cell entry of B.1_pp_ and BA.3_pp_ was Camostat-sensitive and thus TMPRSS2 dependent, although inhibition of BA.3_pp_ entry was slightly less efficient, likely due to the previously noted moderate shift of Omicron subvariants towards cathepsin L for Calu-3 lung cell entry (23, 24). Further, residual Calu-3 lung cell entry driven by the B.1 and BA.3 S proteins lacking the QTQTX motif was at least partially Camostat-resistant, in alignment with the concept that absence of precleavage at the S1/S2 site skews entry towards the cathepsin L-dependent route (5). Finally, Calu-3 cell entry of BA.3.2.2_pp_ was highly Camostat sensitive, irrespective of the deletion of the QTQTX motif, as expected from the efficient precleavage of the BA.3.2.2 S protein at the S1/S2 site.

In sum, we note that the Calu-3 lung cell entry of BA.3 is TMPRSS2-dependent and that protease preference was maintained during BA.3.2 evolution. However, BA.3.2 has apparently lost the need for the QTQTX motif for efficient lung cell entry, suggesting that the flexibility of its cleavage loop does not depend on this motif. It will be interesting to determine whether this is due to mutations in other parts of the S protein or to mutation A688D, the only mutation that distinguishes the BA.3 from the BA.3.2 cleavage loop. It is also noteworthy that the S protein of Omicron subvariant BA.1 exhibits characteristics of being pre-primed for membrane fusion even in the absence of ACE2 binding (25), suggesting reduced requirements for priming that may have been further diminished in the context of the BA.3.2.2 S protein. Finally, we note that roughly 5% of recent XFG sequences harbor mutations in the QTQTX motif, suggesting that also some XFG variants might exhibit reduced dependence on the QTQTX motif.

Collectively, our data suggest that the loss of the first QT repeat in the BA.3.2.2 variant detected in Australia may not have appreciably compromised use of the TMPRSS2-dependent entry route and thus spread in the respiratory epithelium. Further, the deletion did not augment evasion form neutralizing antibodies (**Supplemental Fig. S4-5**). Whether it conferred an advantage to the virus remains to be determined.

## MATERIAL AND METHODS

### Cell culture

All cell lines were maintained at 37 °C in a humidified atmosphere containing 5% CO₂. The cell lines authentication was analysed by short tandem repeat (STR) analysis, using PCR amplification and subsequently sequencing of a cytochrome c oxidase gene fragment, as well as microscopic inspection, and/or cell line-specific growth characteristics. Mycoplasma contamination was examined by regular PCR-based diagnostic. Vero76 cells (African green monkey kidney, female; CRL-1586, ATCC; RRID: CVCL_0574, kindly provided by Andrea Maisner) and 293T cells (human kidney, female; ACC-635, DSMZ; RRID: CVCL_0063) were grown in Dulbecco’s modified Eagle’s medium (DMEM; PAN-Biotech) supplemented with 10% fetal bovine serum (FBS, Biochrom) and 1% penicillin-streptomycin solution (pen/strep; PAN-Biotech). Calu-3 cells (human lung, male; HTB-55, ATCC; RRID: CVCL_0609, kindly provided by Stephan Ludwig) and Calu-3 cells negative for MAVS (Calu-3 KO-MAVS, kindly provided by Caroline Goujon) (19) were maintained in Minimum Essential Medium (MEM) supplemented with 20% FBS, 1% pen/strep, 1% non-essential amino acid solution, and 1 mM sodium pyruvate. A549 cells (human lung, male; CRM-CCL-185, ATCC; RRID:CVCL_0023, kindly provided by Georg Herrler) DMEM/F-12 medium (Thermo Fisher Scientific) supplemented with 10% FBS, 1% pen/strep, 1% non-essential amino acid solution, and 1 mM sodium pyruvate. Transfection of 293T cells was performed by calcium phosphate precipitation.

### Expression plasmids and sequence analysis

Expression plasmids pCAGGS-DsRed(1), pCAGGS-VSV-G (26), pCG1-SARS-CoV-2 B.1 S Δ18 (codon-optimised, C-terminal truncation of 18 amino acid residues) (1), pCG1-SARS-CoV-2 BA.3 SΔ18 (codon-optimised, C-terminal truncation of 18 amino acid residues) (27, 28), pCG1-SARS-CoV-2 BA.3.2.2 S Δ18 (codon-optimised, C-terminal truncation of 18 amino acid residues) (13), pCG1_solACE2-Fc(29), pQCXIP-beta-galactosidase alpha fragment (28), and pQCXIP-beta-galactosidase omega fragment (28) have been described previously while pCAGGS-based plasmids expressing NiV-F or NiV-G were provided by Andre Maisner.

Plasmids encoding SARS-CoV-2 B.1 SΔ18 ΔQT, SARS-CoV-2 B.1 SΔ18 ΔQTQTN, SARS-CoV-2 BA.3 SΔ18 ΔQT, SARS-CoV-2 BA.3 SΔ18 ΔQTQTK, SARS-CoV-2 BA.3.2.2 SΔ18 ΔQT, SARS-CoV-2 BA.3.2.2 SΔ18 ΔQTQTR, were generated from either pCG1-SARS-CoV-2 B.1 SΔ18, pCG1-SARS-CoV-2 BA.3 or pCG1-SARS-CoV-2 BA.3.2.2 (all codon-optimised, C-terminal truncation of 18 residues) by PCR-based introduction of the respective deletions using overlapping primers. All S protein plasmid constructs were sequence-verified by Sanger sequencing (Microsynth SeqLab). The pCG1 expression vector was kindly shared by Roberto Cattaneo (Mayo Clinic, Rochester, MN, USA).

### Pseudovirus particle production

Pseudovirus particles were produced as previously described (30). Briefly, 293T cells were transfected with plasmids encoding the respective S protein, VSV-G, empty vector (EV, negative control) or DsRed (negative control) using calcium phosphate precipitation. The cells were washed with PBS and inoculated with VSV-G-trans-complemented VSV*ΔG (FLuc) (kindly provided by Gert Zimmer) (31) 24 h post transfection. After 1 h of incubation at 37 °C, cells were washed with PBS and further incubated in culture medium supplemented with anti-VSV-G antibody (supernatant from I1-hybridoma cells; ATCC CRL-2700). Cells transfected with VSV-G plasmid received medium without antibody. The supernatants were collected and clarified by centrifugation (4,000 g, 10 min) 16-18 h post-inoculation. Clarified pseudovirus preparations were aliquot and stored at -80 °C until further use.

### Analysis of S protein incorporation into pseudovirus particles

Pseudoviruses carrying the indicated respective S protein were concentrated by centrifugation at 16,800 g for 90 min at 4 °C through a 20% (w/v) sucrose cushion prepared in PBS. The supernatant was removed and pseudoviral pellets were lysed in 2x SDS sample buffer (0.03 M Tris-HCl, 10% glycerol, 2% SDS, 5% β-mercaptoethanol, 0.2% bromophenol blue, 1 mM EDTA) and heated to 96°C for 10 min subsequently. Proteins then were resolved by SDS-PAGE and transferred onto nitrocellulose membranes (Hartenstein). Membranes were blocked for 30 min in PBS-T (PBS with 0.02% Tween-20; Carl Roth) containing 5% skim milk, followed by overnight incubation at 4 °C with primary antibodies against the S2 subunit (rabbit, 1:2000 in PBS-T with 5% skim milk; Biozol, SIN-40590-T62) or the VSV matrix protein (VSV-M, mouse, 1:1000 in PBS-T with 5% skim milk; Kerafast, EB0011). After three washes with PBS-T, membranes were incubated with HRP-conjugated secondary antibodies: anti-rabbit (SARS-2S, 1:2000; Dianova, 111-035-003) or anti-mouse (VSV-M, 1:2000; Dianova, 115-035-003), diluted in PBS-T with 5% skim milk. Blots were washed again three times in PBS-T, and protein bands were detected using a homemade chemiluminescence substrate (0.1 M Tris-HCl [pH 8.6], 250 µg/ml luminol, 0.1 mg/ml para-hydroxycoumaric acid, 0.3% hydrogen peroxide). Signal detection was performed using the Azure 600 imaging system, and data were processed with AzureSpot Pro software (Azure Biosystems).

### Analysis of spike protein-mediated cell entry

To evaluate cell tropism and entry efficiency, equal volumes of pseudotyped VSV particles bearing respective SARS-CoV-2 S proteins, VSV-G (positive control), or lacking glycoprotein (negative control) were inoculated onto target cells seeded in 96-well plates. To assess inhibition by soluble ACE2 or antibodies present in post-vaccination plasma, pseudoviruses were pre-incubated at 37 °C for 30 min with equal volumes of serial dilutions of solACE2-Fc (undiluted, 1:4, 1:16, 1:64, 1:256, 1:1024), serial dilutions of heat-inactivated (30 min at 56 °C) human plasma (1:25, 1:100, 1:400, 1:1600, 1:6400, 1:25600), or with diluent only (culture medium, control) before inoculation. At 16-18 h post-inoculation, viral entry was quantified by measuring firefly luciferase activity encoded by the pseudovirus genome. To assess inhibition by Camostat and MDL28170, target cells were pre-incubated at 37 °C for 60 min with Camostat (25 µM; Sigma-Aldrich) and MDL28170 (12.5 µM; MedChemExpress) or with diluent only (culture medium, control) before inoculation. To assess the effect of Amphotericin B, target cells were pre-incubated at 37 °C for 60 min with Amphotericin B (2.5 µM, Sigma-Aldrich) or with diluent only (culture medium, control) before inoculation. At 16-18 h post-inoculation, viral entry was quantified by measuring firefly luciferase activity encoded by the pseudovirus genome. For the luminescence detection, culture medium was removed and cells were lysed in PBS containing 0.5% Tergitol 15-S-9 (Carl Roth) for 30 min at room temperature (50 µl/well). Cell lysates were transferred into white 96-well plates, mixed with an equal volume of luciferase substrate (Beetle-Juice, PJK), and luminescence was measured using a Hidex Sense luminometer (Hidex).

Neutralization efficiency was determined based on the relative reduction of luciferase activity in plasma-containing samples compared to samples that did not contain plasma (0% neutralisation). Next, dose-response curves were calculated based on a non-linear regression model in order to obtain 50% neutralizing titer (NT50) values.

### Human plasma samples

Post-vaccination plasma samples were obtained from n = 24 individuals two weeks following vaccination with Corminaty LP.8.1. Sampling was performed as part of the COVID-19 Contact (CoCo) Study (German Clinical Trial Registry, DRKS00021152), an ongoing, prospective, observational study monitoring anti-SARS-CoV-2 immunoglobulin G and immune responses in healthcare professionals at Hannover Medical School (32).

### Analysis of spike protein-mediated cell-cell fusion

Cell-cell fusion was analysed as described (1, 17, 28, 33, 34). Briefly, effector 293T cells were transfected with beta-galactosidase alpha fragment expression plasmid jointly with plasmids coding for the respective S protein, Nipah virus fusion and glycoprotein (NiV-F, NiV-G; positive control), or empty vector (negative control). In addition, target 293T cells were co-transfected with plasmid encoding for the beta-galactosidase omega fragment jointly with human ACE2 expression alone or in combination with TMPRSS2 expression plasmid. Following 18 h of incubation, both effector and target cells were washed with PBS and maintained in fresh medium for 6 h. Target cells were subsequently resuspended in fresh medium and added onto effector cells. After 18 h of co-culture, Gal-Screen substrate (Thermo Fisher Scientific) was applied, and luminescence was measured 90 min later with a Hidex Sense luminometer.

### Protein models and sequence analysis

Protein models for trimeric S protein ectodomains and S1/S2 loops for the SARS-CoV-2 B.1 (GISAID Accession identifier: EPI_ISL_425259), BA.3 (EPI_ISL_8801154) and BA.3.2.2 (EPI_ISL_19893762) lineages were constructed by homology modeling with the crystal structure PDB:6XR8 (35) using the SWISS-MODEL tool (https://swissmodel.expasy.org/). Structural analysis, visualization and coloring was further done using UCSF Chimera (version 1.17.3) software (36). For alignments, S1/S2 loop sequences from the following S proteins were used: hCoV-19/Wuhan/Hu-1/2019 (GISAID: EPI_ISL_402125), hCoV-19/South Africa/NICD-N25677/2021 (GISAID: EPI_ISL_8801154), hCoV-19/Germany/NW-RKI-I-1148174/2025 (EPI_ISL_19893762), hCoV-19/bat/Laos/BANAL-52/2020 (GISAID: EPI_ISL_430244), hCoV-19/bat/Laos/BANAL-103/2020 (GISAID: EPI_ISL_4302645), hCoV-19/bat/Laos/BANAL-236/2020 (GISAID: EPI_ISL_4302647), hCoV-19/bat/Yunnan/RaTG13/2013 (GISAID: EPI_ISL_402131), Bat SARSr-CoV BtSY2 Bat/2018/S18CXBatR24 (GenBank: WBV74286.1), hCoV-19/bat/Cambodia/RShSTT182/2010 (GISAID: EPI_ISL_852604), hCoV-19/bat/Cambodia/RShSTT200/2010 (GISAID: EPI_ISL_852605), hCoV-19/pangolin/Guangdong/1/2019 (GISAID: EPI_ISL_410721), hCoV-19/pangolin/Guangdong/A22-2/2019 (GISAID: EPI_ISL_471467), Pangolin SARSr-CoV MP789 (GenBank: QIG55945.1),

hCoV-19/pangolin/Guangdong/FM45-9/2019 (GISAID: EPI_ISL_471468), hCoV-

19/pangolin/Guangdong/SM44-9/2019 (GISAID: EPI_ISL_471469), hCoV-

19/pangolin/Guangdong/SM79-9/2019 (GISAID: EPI_ISL_471470), hCoV-

19/pangolin/Guangdong/cDNA8-S/2019 (GISAID: EPI_ISL_471461), hCoV-

19/pangolin/Guangdong/cDNA9-S/2019 (GISAID: EPI_ISL_471462), hCoV-

19/pangolin/Guangdong/cDNA16-S/2019 (GISAID: EPI_ISL_471463), hCoV-

19/pangolin/Guangdong/cDNA18-S/2019 (GISAID: EPI_ISL_471464), hCoV-

19/pangolin/Guangdong/cDNA20-S/2019 (GISAID: EPI_ISL_471465), hCoV-

19/pangolin/Guangdong/cDNA31-S/2019 (GISAID: EPI_ISL_471466), hCoV-

19/pangolin/Guangxi/P1E/2017 (GISAID: EPI_ISL_410539), hCoV-

19/pangolin/Guangxi/P2V/2017 (GISAID: EPI_ISL_410542), hCoV-

19/pangolin/Guangxi/P4L/2017 (GISAID: EPI_ISL_410538), hCoV-

19/pangolin/Guangxi/P5E/2017 (GISAID: EPI_ISL_410541), hCoV-

19/pangolin/Guangxi/P5L/2017 (GISAID: EPI_ISL_410540), Bat SARSr-CoV Rc-mk2

(GenBank: BDD37268.1), Bat SARSr-CoV Rc-o319 (GenBank: LC556375.1), and Bat SARSr-CoV Rc-os20 (GenBank: BDD37258.1). Sequence alignments were done using the Multiple Sequence Alignment by CLUSTALW web tool (https://www.genome.jp/tools-bin/clustalw) with standard parameters.

### Data analysis

Data analysis was conducted using Microsoft Excel (Office Professional Plus 2016; Microsoft Corporation) and GraphPad Prism version 10.6.0 (GraphPad Software). Statistical significance was determined using one-or two-way analysis of variance (ANOVA) with Tukey’s multiple comparison test, or Kruskal-Wallis test with Dunn’s multiple comparisons test (the exact test is stated in the figure legends). P values ≤0.05 were considered statistically significant (ns, p >0.05; *, p ≤0.05; **, p ≤0.01; ***, p ≤0.001).

## Acknowledgements

We acknowledge the originating laboratories responsible for sample collection and the submitting laboratories that generated and shared genome data via GISAID, which formed the basis of this study. We also thank Roberto Cattaneo, Caroline Goujon, Georg Herrler, Stephan Ludwig, Andrea Maisner and Gert Zimmer for providing reagents. We also thank the participants of the CoCo Study for their contribution, as well as the entire CoCo Study team for their continued support. Calu-3 KO-MAVS, kindly provided by Caroline Goujon, CNRS, University of Montpellier, France. S.P. acknowledges funding by the EU project UNDINE (grant agreement number 101057100), the COVID-19-Research Network Lower Saxony (COFONI) through funding from the Ministry of Science and Culture of Lower Saxony in Germany (14-76403-184, projects 7FF22, 6FF22, 10FF22), the European Commission, Horizon Europe, VIGILANT (101041799) and the German Center for Infection Research (TTU 01.829).

